# Improved two-step testing of genome-wide gene-environment interactions

**DOI:** 10.1101/2022.06.14.496154

**Authors:** Eric S. Kawaguchi, Andre E. Kim, Juan Pablo Lewinger, W. James Gauderman

**Affiliations:** Department of Population and Public Health Sciences, University of Southern California

## Abstract

Two-step tests for gene-environment (*G* × *E*) interactions exploit marginal SNP effects to improve the power of a genome-wide interaction scan (GWIS). They combine a screening step based on marginal effects used to ‘bin’ SNPs for weighted hypothesis testing in the second step to deliver greater power over single-step tests while preserving the genome-wide type I error. However, the presence of many SNPs with detectable marginal effects on the trait of interest can reduce power by ‘displacing’ true interactions with weaker marginal effects and by adding to the number of tests that need to be corrected for multiple testing. We introduce a new significance-based allocation into bins for step 2 *G* × *E* testing that overcomes the displacement issue and propose a computationally efficient approach to account for multiple testing within bins. Simulation results demonstrate that these simple improvements can provide substantially greater power than current methods under several scenarios. An application to a multi-study collaboration for understanding colorectal cancer (CRC) reveals a *G*×Sex interaction located within the SMAD7 gene.

## 1 Introduction

Identifying gene-environment interactions (*G* × *E*) is critical for understanding how health is affected by both an individual’s genetic background (*G*) and exposure to environmental factors (*E*). Genome-wide interaction scans (GWIS) test *G* × *E* interactions one-SNP-at-a-time by modeling the genotype, an environmental exposure, and the corresponding interaction term. Testing of the *G* × *E* is based on the significance of the interaction term, typically from a logistic (for a binary/disease trait), linear (for a quantitative trait), or Cox (1972) (for a survival trait) regression model. A standard one-step GWIS proceeds by testing each *G* × *E* interaction at significance level of *α*^*^ = 5 × 10^-8^, common in genome-wide association studies (GWAS) or GWIS studies (Dudbridge and Gusnanto, 2008). However, the statistical power to detect an interaction in a one-step GWIS is generally much lower than the power for detecting a genetic marginal effect in a GWAS.

Two-step tests for GWIS have been proposed to improve the power of a *G* × *E* analysis while controlling the FWER for disease (Kooperberg and LeBlanc, 2008; Murcray et al., 2009, 2011; Hsu et al., 2012; Gauderman et al., 2013; Wang et al., 2021), quantitative traits (Paré et al., 2010; Zhang et al., 2016), and time-to-event traits (Kawaguchi et al., 2022). In all of these two-step procedures, independent information on *G* × *E* not captured by the standard *G* × *E* test, is used to perform an initial screening (Step 1) to prioritize SNPs that are more likely to be involved in an interaction. These SNPs are formally tested for an interaction (Step 2) under a modified *α*^*^, thus reducing the multiple testing burden (Kooperberg and LeBlanc, 2008; Murcray et al., 2009).

The marginal outcome-gene association statistic derived from modeling the outcome on each gene individually is a commonly-used screening statistic for quantitative (Zhang et al., 2016), binary/disease (Kooperberg and LeBlanc, 2008), and time-to-event (Kawaguchi et al., 2022) traits. For case-control studies the exposure-gene association statistic, modeling the relationship between each gene on the exposure, can also be informative (Murcray et al., 2009). Methods that utilize both outcome-gene and exposure-gene associations in a case-control study have also been developed (Murcray et al., 2011; Hsu et al., 2012; Gauderman et al., 2013). A key requirement for validity of any two-step procedure is that the statistics used in Step 1 and Step 2 are asymptotically independent (Dai et al., 2012; Kawaguchi et al., 2022).

There are two widely-used procedures for prioritizing SNPs in Step-2 *G* × *E* testing after the Step-1 screening: subset (Kooperberg and LeBlanc, 2008; Murcray et al., 2009) and weighted hypothesis testing (Ionita-Laza et al., 2007). In subset testing of the *M* total SNPs that are being scanned, only the *m << M* SNPs that pass a significance threshold based on the screening statistic are tested in Step 2 using a standard *G* × *E* test. The significance threshold in step 2 for *G* × *E* discovery is calculated using a Bonferroni correction that is based on the number of SNPs that pass the screening (*α*^*^ = *α/m*), which is much less stringent than the threshold used in a single step approach. A trade off for a relaxed threshold is that SNPs that do not pass the step 1 screening will not be tested. An alternative approach that does not rely on a pass/no pass hard rule is weighted hypothesis testing. Here, SNPs are allocated into bins based on the magnitude of the screening statistic. Each bin has a corresponding bin-wise error rate (BWER) such that the sum across all bins does not exceed *α*. Lower bins are allocated a larger fraction of *α* (see Section 2.1 for more detail), so that SNPs in those bins are tested at a more liberal significance threshold. Conversely, SNPs that are placed in higher bins are tested at a much more stringent BWER. Unlike subset testing, every SNP is tested in Step 2 of the weighed approach; yet SNPs that are more likely to have an interaction based on the screening statistic will have a higher chance of being discovered. Although weighted hypothesis testing is often more powerful than subset testing (Ionita-Laza et al., 2007; Gauderman et al., 2013), there is no universally most powerful approach.

The motivation behind two-step hypothesis testing is that in the presence of a true *G* × *E* interaction effect, one can typically expect there to be marginal *G* effects on the outcome, which makes the marginal outcome-gene statistic useful for screening/ranking SNPs in Step 1 (e.g being placed in bin 1, the bin with the largest BWER). However, this only proves useful if not too many SNPs have sizeable marginal effects but no *G* × *E* interaction. This is often not the case in a GWIS where known “hits” from a prior GWAS provides a set of SNPs for which there is a strong marginal outcome-gene effect. For example, more than 140 GWAS-significant (marginal-effect) loci have been previously identified for colorectal cancer (Huyghe et al., 2019) and the majority of these likely do not exhibit a *G* × *E* interaction. This will result in a phenomena we refer to as “bin overcrowding”, where these SNPs will overcrowd the earlier bins in the weighted testing approach due to having non-zero marginal effects. Thus, even if a true *G* × *E* effect induces a non-zero marginal effect, it is competing against other known (or previously unknown) non-zero marginal effects which can force the true *G* × *E* effect to not be optimally tested (e.g. being placed/tested in a later bin with a stricter BWER). To avoid this loss of power, one can filter out these loci in advance. However, this approach requires prior knowledge of loci with marginal effects, and it also removes these SNPs from consideration for *G* × *E* testing.

The contribution of this paper is the improvement of two-step GWIS testing in two ways. First, we propose a simple yet effective approach to prioritize tests in the screening step that minimizes the potential power loss due to the presence of SNPs with a marginal effect but no interaction effect. Second, we show how additional accounting for correlation among SNPs in linkage disequilibrium (LD) in the testing step can further increase power. We demonstrate via simulation that this new two-step testing method yields greater power than its predecessors. We apply the approach to identify *G* × *E* interactions for colorectal cancer.

We describe current methods for testing *G* × *E* interactions using the two-step hypothesis testing paradigm and our proposed approach in Section 2. Simulation studies that compare the performance of two-step hypothesis testing procedures are given in Section 3 and an application to a GWIS for colorectal cancer is provided in Section 4. Lastly, in Section 5, we provide concluding remarks, limitations, and areas of future research.

## 2 Methods

Consider a gene-environment interactions study with a (continuous, binary, or time-to-event) trait/outcome *Y*, an environmental exposure of interest *E*, and *M* SNPs (*G_j_, j* = 1,…, *M*) measured or imputed for each of the *N* subjects. To perform a GWIS, we assume *M* tests of *G* × *E* interaction with test statistics 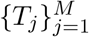 and corresponding *p*-values 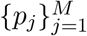 are computed. If for example the trait *Y* is quantitative, a standard one-step GWIS generates the *G* × *E* interaction one-at-a-time by using the following model:

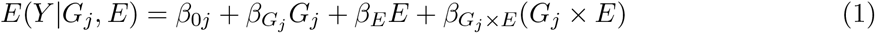

for each of the *M* SNPs. For ease of exposition, we did not include subject-level covariates in the model, but in practice adjustment covariates like sex, age, and principal components capturing genetic ancestry are critical for preventing spurious associations. Each *T_j_* corresponds to the test statistic for testing the null hypothesis *H*_0_: *β_G_j_ ×E_* = 0. An adjustment for multiple comparisons is applied to preserve the family-wise Type I error rate (FWER) at a prespecified significance level *α* (e.g., *α*^*^ = 5 × 10^-8^ or *α*^*^ = *α/M*). However, this correction will lead to low power in detecting a *G* × *E* interaction.

### 2.1 Two-step tests

Two-step methods have been developed to improve the power of GWIS(Kooperberg and LeBlanc, 2008; Murcray et al., 2009; Ionita-Laza et al., 2007). The test of interaction is also based on *T_j_* (Step 2); however, the significance level at which each of the *M* SNPs is tested depends on an initial screening step (Step 1) based on a screening statistics *S_j_* computed for each SNP. A common choice of screening statistic is based on the marginal effect of a SNP on the trait, as the presence of an interaction will typically induce a marginal *G* effect on the outcome (Kooperberg and LeBlanc, 2008). For a quantitative trait, the marginal outcome-gene association can be modeled using

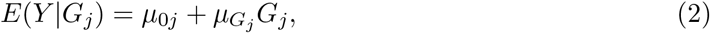

and *S_j_* is the test statistic corresponding to *H*_0_: *μ_G_j__* = 0. Model 2 has the form typically used to identify SNPs associated with the outcome in a standard GWAS.

Two approaches to two-step testing are being widely used: subset testing (Kooperberg and LeBlanc, 2008) and weighted hypothesis testing (Ionita-Laza et al., 2007). <u>Subset testing</u>: In subset testing, each of the *M* screening statistics is compared to a prespecified significance threshold *α*_0_. Let 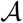 be the collection (subset) of indices of SNPs that pass the Step 1 screen (i.e. SNPs for which *S_j_* is statistically significant at the *α*_0_ level) with 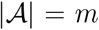, where |*A*| represents the cardinality of set *A*. Then, for each 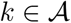, the test statistic *p*-value *p_k_* is compared against the Bonferroni-corrected significance level *α*^*^ = *α/m*, which is a less stringent significance level than in the standard single-step GWIS. Note that tests where 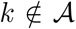 will never be tested – or equivalently tested against a significance level of *α*^*^ = 0. This ensures that the overall Type I error rate is retained at *α*. <u>Weighted testing</u>: Instead of only testing a subset of hypotheses based on some dichotomous screening procedure, one can test all *M* test statistics in Step 2 using a weighted hypothesis test (Ionita-Laza et al., 2007) where the significance level assigned to each Step 2 SNP is based on the ordered (largest to smallest) absolute values of 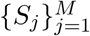 from Step 1. The rationale behind the weighted screening approach is that more promising SNPs - as measured by the screening statistic – are tested at a less stringent significance level. SNPs are assigned into *B* bins according to their *M* screening statistics. To control the FWER at level *α*, we derive bin-wise error rates (BWER) such that *α*_1_ + *α*_2_ +… + *α_B_* ≤ *α*. A common choice is *α_b_* = *α*/2^*b*+1^; then, for sufficiently large *B*,

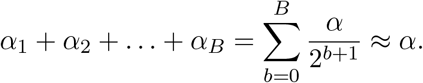

By partitioning *α* in such a way, the overall Type I error rate is controlled at *α* while allocating a greater fraction of *α* to the top bins (i.e. those with the most promising SNPs). A Bonferroni-like correction can be used to preserve BWER by dividing *α_b_* by 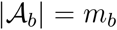, where 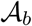 is the set of indices for tests in bin *b* (size of the bin). We are now left with the non-trivial task of deciding how many SNPs should be allocated to each bin. Using a predetermined initial bin size, *B*_0_, Ionita-Laza et al. (2007) suggested binning tests such that *m_b_* = 2^*b*–1^*B*_0_. For example, the SNPs with the *B*_0_ largest values of |*S_j_*| are placed in 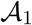 and tested against a significance threshold of *α*_1_/*B*_0_, the next 2B0 SNPs are tested in bin 2 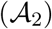 at level *α*_2_/2*B*_0_ and so forth. Ionita-Laza et al. (2007) suggested setting *B*_0_ = 5 so that *m*_1_ = 5, *m*_2_ = 10, *m*_3_ = 20, *m*_4_ = 40… Since the binning of SNPs can be done in several ways, we refer to the Ionita-Laza et al. (2007) approach as *rank-based* (*RB*) binning to contrast with significance-based binning that we propose below. Two-step RB-weighted tests are in general, more powerful than subset testing (Gauderman et al., 2013).

### 2.2 Proposal # 1: Significance-based (SB) allocation of SNPs to bins in Step 1

Note that subset testing can be seen as a unique case of RB-weighted testing with two bins with *α*_1_ = *α*, 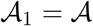, *α*_2_ = 0, and 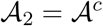 where 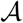 is the subset of indices of SNPs that have a Step 1 *p*-value < *α*_0_. Unlike RB-weighted testing, where the size of bin 1 is set *a priori*, here the size of the bin is determined by the number of *p*-values in Step 1 that fall within the interval (0, *α*_0_). Rather than creating two bins based off of one *p*-value (e.g. *α*_0_), one can allocate tests into *B* bins using a series of significance-level cutoffs. More specifically, defining *τ* = (*τ*_1_, *τ*_2_,…, *τ*_*B*+1_) as the set of significance level cutoffs, the collection of tests in bin *b* is 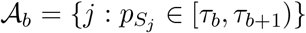. We refer to this type of screening as significance-based (SB) weighted testing, since SNP prioritization is based on the significance levels of the *p*-values corresponding to each screening statistic (*p_S_j__*). Thus, SB-weighted testing can be viewed as a hybrid between both subset and RB-weighted testing. While *τ* can be defined arbitrarily, we propose to set *τ* = (0, *B*_0_/*M*, 3*B*_0_/*M*, 7*B*_0_/*M*,…, 1) which, in expectation, corresponds to binnings that are identical to RB-weighted testing. However, bin sizes are not capped since *p*-values are used to determine bin allocation.

Assuming that the null hypothesis of no marginal association holds for all *M* SNPs, each *p_S_j__* is uniformly distributed in the interval [0, 1]. In expectation, Bo of the marginal outcome-gene screening statistics should have a *p*-value less than *B*_0_/*M*, 2*B*_0_ of them should be in the interval (*B*_0_/*M*, 3*B*_0_/*M*), and so forth. However, this relationship only holds under the null hypothesis of no outcome-gene associations. In practice, this assumption may not hold and will lead to a more-than-expected number of *p*-values to lie in the top bins. This overcrowding of the top bins (e.g., 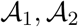) will reduce the power of the RB-weighted testing approach since bin sizes are capped and SNPs with true *G* × *E* effects may be pushed to a bin in which the interaction will be tested using a stricter threshold. As we will show in our numerical studies, the performance of RB-weighted testing is dependent on both the strength and number of non-null outcome-gene associations, which typically is unknown *a priori*. While overcrowding is still an issue with SB-weighted testing, we will show that the downstream effect on the testing step is reduced compared to RB-weighted testing.

### 2.3 Proposal #2: Accounting for correlation among tests in Step 2

While two-step hypothesis testing generally provides increased power over the standard one-step GWIS, a Bonferroni-type correction is still used to control the FWER in Step 2. This adjustment tends to be overly conservative in situations where correlation is present as it is the case in association studies due to linkage disequilibrium (LD). Accounting for genetic correlation can provide additional power gains.

Permutation tests are seen as the gold standard by permuting the data in a way that simulates the null hypothesis while simultaneously maintaining the original correlation structure. However, in large association studies, permutation-based approaches are computationally prohibitive. Furthermore, it is not obvious how to extend permutation tests in the *G* × *E* setting due to the hierarchical structure of the model. Several more efficient methods to account for correlation between tests have been proposed. For example, Conneely and Boehnke (2007) developed a method (*p_ACT_*) that attains the accuracy of permutation or simulation-based tests in much less computation time. Their approach, however, requires valid estimation of the covariance matrix, which is computationally prohibitive if applied genome wide. Another alternative is to replace the denominator of the Bonferroni correction, the total number of tests within a bin, with an estimate of the effective number of independent tests (*M*^*^). Cheverud et al. (1983), among others, have shown that the overall correlation among variables in a set can be captured by the variance of the eigenvalues derived from their correlation matrix (i.e. higher correlation among variables will lead to higher eigenvalue variance).

Let 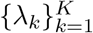 be the set of eigenvalues obtained by a principal component analysis (PCA) from the pair-wise SNP correlation matrix (i.e. LD matrix), arranged in decreasing order. Gao et al. (2008) proposed a simple and fast correction (simple*M*) that well approximates permutation-based corrections in both simulated and real data based on these eigenvalues

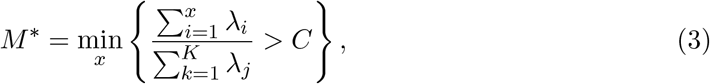

where *C* determines the percentage cutoff of explained variation. Gao et al. (2008) suggest setting *C* = 0.995 so that *M*^*^ corresponds to the minimum number of eigenvalues needed to explain at least 99.5% of the variation in the data.

Applying this correction genome wide is infeasible as it requires, first, the calculation of an *M* × *M* correlation matrix and, second, an eigendecomposition (e.g. PCA) to derive the eigenvalues. We instead apply the correction to the set of SNPs within each bin. For example, if we let **G** be the *N* × *M* matrix of SNP genotypes (or imputed dosages) for *N* subjects, then we calculate 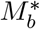 for bin *b* using the correlation matrix of 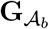, the submatrix of **G** that corresponds to the indices in 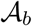. Note that, by design, 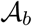 grows larger, in expectation, as *b* increases. Thus the calculation of *M*^*^ for the larger bins may still be computationally demanding. In practice, as shown in Section 3, we restrict the computation of 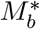 for the first seven bins and set the significance threshold for the later bins to be 0 since the threshold to be declared statistically significant significant in those upper bins is extremely stringent.

## 3 Simulation studies

We compare the performance of our two-step significance-based (SB) weighted testing procedure to 1) the standard one-step GWIS, and 2) the two-step rank-based (RB) weighted testing procedure. For both two-step RB and SB-testing, we compare the standard Bonferroni correction to the LD-adjusted Bonferroni correction using the simple*M* approach (Gao et al., 2008) to account for LD within bins. As recommended by Gao et al. (2008), we set the tuning parameter *C* = 0.995.

Let **G** be an *N* × *M* genotype matrix for *N* individuals and *M* SNPs. In all of our simulations, we assume *M* = 25, 000. We partition the *M* = 25, 000 SNPs into blocks of 50 SNPs such that **G** = [**G**_1_, **G**_2_,…, **G**_500_], where ***G**_j_* is the *j*th block of *N* × 50 SNPs. Each ***G**_j_* is simulated based on sampled minor allele frequencies (MAFs) and LD-matrices from the 1000 Genomes Project. For clarity, we denote *G_j_* as the *j*th SNP and **G***_j_* as the *j*th block. Quantitative traits are simulated according to the following linear model:

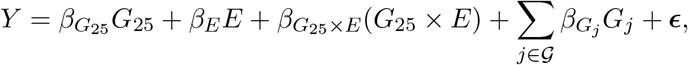

where 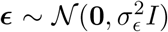 for some 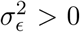, *E* is the exposure variable (assumed to be binary) with Pr(E = 1) = 0.3 and 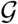 corresponds to the set of SNPs that are only marginally associated with the outcome but have no *G* × *E* effect (G-only loci). By construction, the 25th SNP within block 1 (**G**_1_) has a true *G* × *E* effect on the outcome (i.e. the *G* × *E* locus). We refer readers to the Appendix for more information on the construction of **G** and the simulation setup.

The value of the parameters *β*_*G*_25__, *β_E_*, and *β*_*G*_25_×*E*_ were set to achieve a predetermined *R*^2^ for each term: 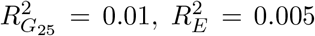 and 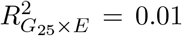. Given a minor allele frequency of 0.231 for the *G* × *E* locus and setting *V ar*(*Y*) = 1, yields *β*_*G*2^5^_ ≈ 0.06, *β_E_* ≈ −0.01 and *β*_*G*25×*E*_ ≈ 0.37. Each *G*-only loci was placed in a different block (i.e. 25th SNP in **G**_2_, 25th SNP in **G**_3_, etc.) so that these SNPs, and SNPs in LD, are expected to be prioritized in Step 1 and thus potentially affect power in identifying the true *G* × *E* effect due to bin overcrowding. For simplicity, we set the expected *R*^2^ 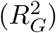 for each of the *G*-only effects to be the same and vary the number of *G*-only effects (i.e. 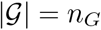).

The testing statistics, *T_j_*, are based on the hypotheses *H*_0_: *β_G_j×E__* = 0 for *j* = 1,…, *M*, where *β_G_j_×E_* is estimated based on the following one-SNP-at-a-time model

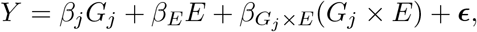

where *G_j_* denotes the *j*th SNP (*j* = 1,…, *M*). For both two-step procedures (RB and SB-weighted), the screening statistics used in Step 1 are based on the test statistics corresponding to the marginal outcome-gene association (Kooperberg and LeBlanc, 2008) based on

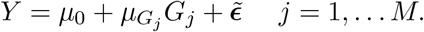

The initial bin size *B*_0_ was set to 5 SNPs for the RB-weighted approach and to an expectation of 5 SNPs assuming no genetic effects for the SB-weighted approach. For the latter, this corresponds to *τ* = (0, 5/25000, 15/25000, 35/25000,…, 1). Under this scheme, we expect both RB and SB-weighted hypothesis testing to have comparable performance for detecting *G* × *E* loci when *G*-only effects are weak or absent. The FWER was set to *α* = 0.05. As aforementioned, calculation of the effective number of independent tests for both two-step approaches will be computationally demanding for bins with larger number of SNPs. To avoid this, we restrict computation of *M*^*^ to the first seven bins and set the significance threshold in the later bins to 0.

Since SNPs in **G**_1_, the block with the causal *G* × *E* locus, are correlated, power is calculated as the number of times we reject the null hypothesis for any of the SNPs in **G**_1_ at the corresponding significance level. FWER is defined as rejecting the null hypothesis *H*_0_: *β_G_j_×E_* = 0 for *j* = 51,…, 25000 at the corresponding significance level. Furthermore, we recorded the ranking of the *G* × *E* locus in terms of its Step 1 statistic as well as the smallest ranking of any loci in **G**_1_ in terms of its Step 1 statistic. Results are averaged over 5,000 Monte Carlo replications.

In our first set of simulations, we set *N* = 2, 000 and 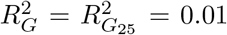 so that the marginal effects of the *G* × *E* locus and *G*-only loci are comparable. Power to detect the true *G* × *E* effect *β_G_j_×E_* with the standard one-step GWIS is approximately 45% (Figure 1 Panel A). When no *G*-only effects are present (*n_G_* = 0), both approaches (RB and SB) have comparable power substantially higher than the standard one-step GWIS. The comparability of RB and SB-weighted testing in this scenario is expected since the binning of tests for both approaches should be nearly identical with no additional *G*-only effects. This is further supported by Figure 2 Panel A, which shows the distribution of the bin placement of the *G* × *E* locus. However, power for RB-weighted testing is sensitive to the number of *G*-only effects, dropping from ≈ 85% when only *n_G_* = 10 *G*-only effects are present to ≈ 64% when *n_G_* = 80 are present (Figure 1). This decrease in power is expected since the *G*-only loci, and their LD-regions, are being ranked higher than the *G* × *E* loci, and hence pushing the *G* × *E* loci into bins with more stringent significance thresholds in the testing step (Figure 2 Panel B). The loss in power for SB-weighted testing is less dramatic, which suggests that it is more robust to bin flooding than RB-weighted testing. Furthermore, adjusting for LD-based correlations in Step 2 provides a consistent increase in power across all scenarios (Figure 1).

**Figure 1:**
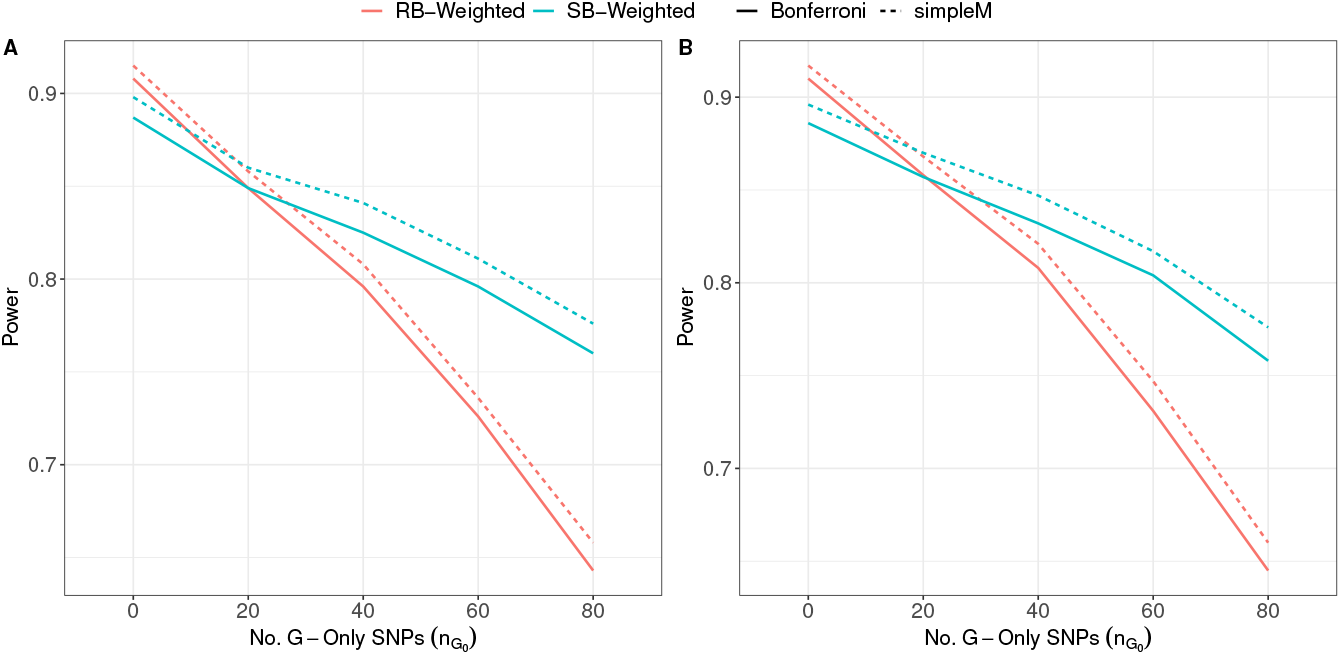
Estimated power when *n_G_ G*-only effects are present (*n_G_* ∈ {10, 20, 40, 80}) that each explain 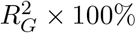 of the variation. RB-Weighted: Rank-based weighted hypothesis testing using with initial bin size *B*_0_ in Step 1; SB-Weighted: Significance-based weighted hypothesis testing using *τ* = (0, *B*_0_/25000, 3*B*_0_/25000,…, 1) as the *p*-value cutoffs in Step 1. Bonferroni: Standard Bonferroni correction within bin; simple*M*: The simple*M* procedure proposed by Gao et al. (2008) with *C* = 0.995. Results are averaged over 5,000 simulations. Panel A: 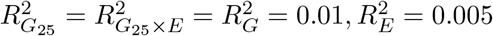, *N* = 2, 000; Panel B: 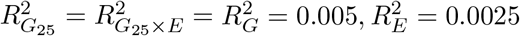, *N* = 4, 000.

**Figure 2:**
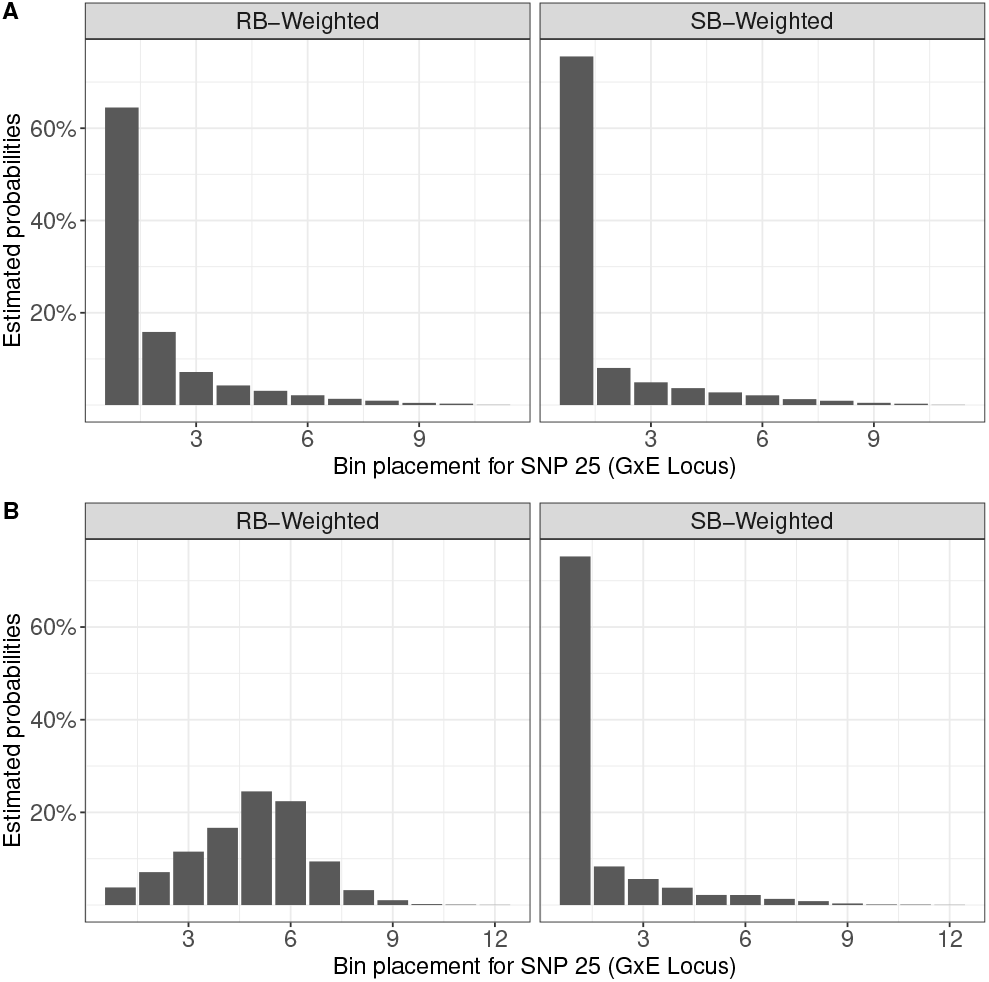
Bar chart of bin placement for the 25th SNP (i.e. *G* × *E* locus) in Step 1 over 5,000 simulations. RB-Weighted: Rank-based weighted hypothesis testing using with initial bin size *B*_0_ in Step 1; SB-Weighted: Significance-based weighted hypothesis testing using *τ* = (0, *B*_0_/25000, 3*B*_0_/25000,…, 1) as the *p*-value cutoffs in Step 1. Simulation parameters: 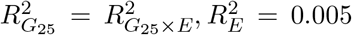, *N* = 2, 000, *M* = 25, 000. Panel A) *n_G_* = 0 *G*-only SNPs; Panel B: *n_G_* = 80 *G*-only SNPs with 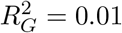.

In a second set of simulations (Figure 1 Panel B) we doubled the sample size *N* to 4,000 and halve the expected *R*^2^ for each factor (i.e. 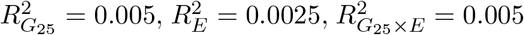, and 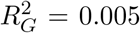). Our results are consistent to what we have seen above which suggests that the power benefit of SB-weighted testing, coupled with the simple*M* procedure, can be achieved under more realistic effect sizes to what is commonly detected in GWAS and GWIS studies. As shown in Figure S1, the overall FWER is preserved at < 5% for the RB and SB approaches in all of the above simulation scenarios.

In addition to increasing the number of marginal effects, we also evaluate both weighted testing methods when we vary both the size and magnitude of the *G*-only SNPs (Table 1). In this set of experiments, we fixed the % of explained variation explained by the *G*-only SNPs to 40% and vary *n_G_* from 10, 20, 40 to 80 such that the expected *R*^2^ for each *G*-only SNP is 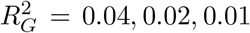 and 0.005, respectively. We keep 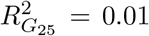 for the causal *G* × *E* locus so that we capture scenarios where the ‘competition’ in prioritizing the causal locus ranges from weak to strong. When 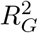 is large relative to 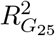, the test for the causal locus has little-to-no chance of being in the top bins and will thus always be tested at a more stringent significance threshold. Power for the standard one-step GWIS is unaffected by the magnitude and number of the *G*-only SNPs. The difference in power is largest between the SB and RB-weighted testing methods when a few number of ‘strong’ *G*-only SNPs (*n_G_* = 10, 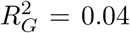) are present (median rank of the Step 1 statistic for the *G* × *E* locus = 61). In this scenario, RB-weighted testing has around 70% power of detecting the *G* × *E* locus whereas SB-weighted testing has power closer to 80%. On the other hand, power between both RB and SB-weighted testing are comparable when more but weaker *G*-only SNPs are present (around 80% power when *n_G_* = 80 and 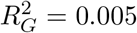). Thus, both magnitude of effect and the number of *G*-only loci can greatly affect power for RB-weighted hypothesis testing. Conversely, SB-weighted screening is more robust to the size and magnitude of the *G*-only SNPs, with power hovering around 80 – 81% in all four scenarios. This is also reflected in the bin placement of the simulated true *G* × *E* loci (Figure S2). The overall FWER is also preserved at < 5% for the RB and SB approaches across these additional scenarios. (Table S1).

**Table 1:** Estimated power when *n_G_ G*-only effects are present (*n_G_* ∈ {10, 20, 40, 80}) and the total amount of variation explained is fixed at 40% 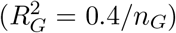. One-step GWIS: The standard one-step GWIS. RB-Weighted: Rank-based weighted hypothesis testing proposed by Ionita-Laza et al. (2007) with *B*_0_ = 5; SB-Weighted: Our proposed significance-based weighted hypothesis testing with *τ* = (0, 5/25000, 15/25000,…, 1) as the *p*-value cutoffs. Bonferroni: Standard Bonferroni correction within bin; simple*M*: The simple*M* procedure proposed by Gao et al. (2008) with *C* = 0.995. Results are averaged over 5000 simulations.

|  |  |  |  |  |
| --- | --- | --- | --- | --- |
| $n_G =$ | 10 | 20 | 40 | 80 |
| $R_G^2 =$ | 0.04 | 0.02 | 0.01 | 0.005 |
| **One-step GWIS** | 0.439 | 0.437 | 0.443 | 0.442 |
| **RB-Weighted** |  |  |  |  |
| Bonferroni | 0.694 | 0.670 | 0.726 | 0.801 |
| simpleM | 0.713 | 0.688 | 0.736 | 0.812 |
| **SB-Weighted** |  |  |  |  |
| Bonferroni | 0.814 | 0.797 | 0.796 | 0.812 |
| simpleM | 0.829 | 0.811 | 0.812 | 0.826 |

It has been shown that the choice of *B*_0_ affects power for weighted testing (Ionita-Laza et al., 2007; Lewinger et al., 2013). We investigate the performance of both RB and SB-weighted hypothesis testing under various initial bin sizes (Figure 3). For this simulation, we assume 80 *G*-only SNPs with 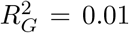 are present (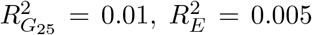 and 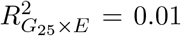) with *N* = 2, 000 and *M* = 25, 000. We see that RB-weighted screening is sensitive to *B*_0_, with an increase in power as *B*_0_ increases. The increase in power can be explained by the fact that we allow more tests to be included in each bin. Thus, even when comparably-sized *G*-only effects are competing with the causal *G* × *E* locus for bin placement, the probability of being in a smaller bin, and thus tested against a less stringent BWER, is higher. However, as others have explored, power can be sensitive to the choice of *B*_0_, and larger values of *B*_0_ can correspond to lower power when no *G*-only effects are present(Lewinger et al., 2013; Gauderman et al., 2013). In contrast, SB-weighted screening is relatively robust to the choice of *B*_0_ used in creating the significance thresholds.

**Figure 3:**
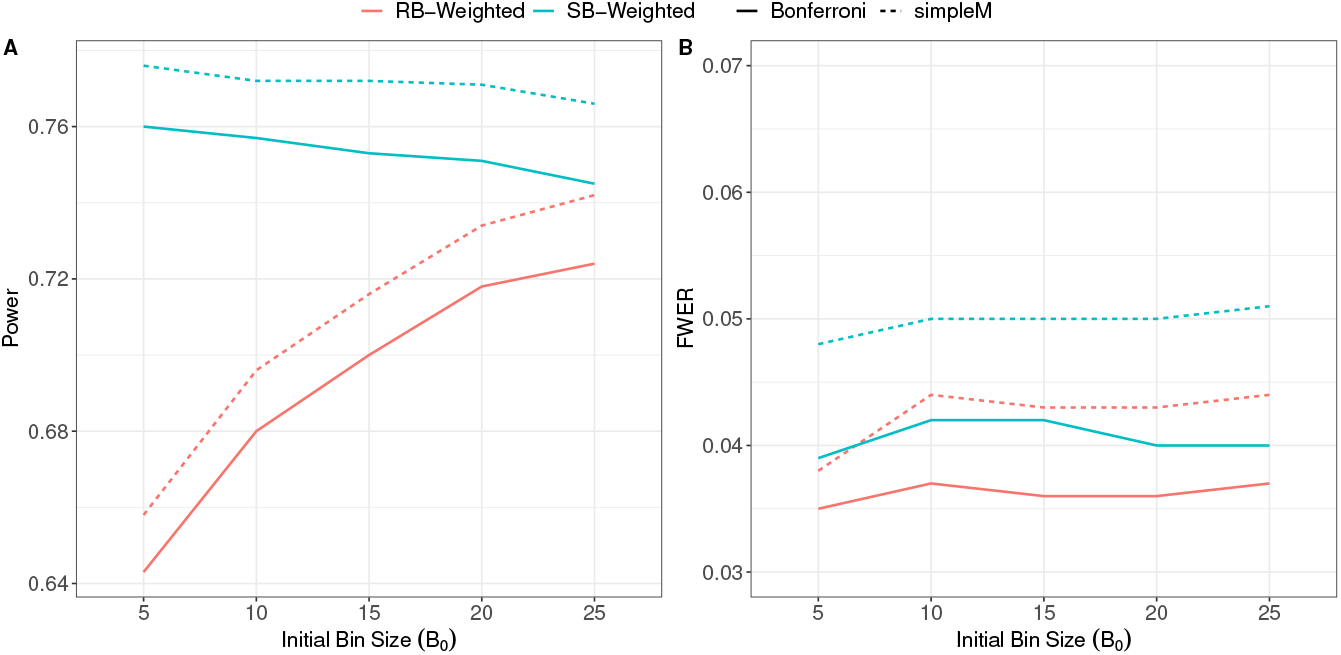
Estimated power and FWER as a function of the initial bin size *B*_0_. 80 *G*-only effects are present with 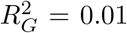. RB-Weighted: Rank-based weighted hypothesis testing using with initial bin size *B*_0_ in Step 1; SB-Weighted: Significance-based weighted hypothesis testing using *τ* = (0, *B*_0_/25000, 3*B*_0_/25000,…, 1) as the *p*-value cutoffs in Step 1. Bonferroni: Standard Bonferroni correction within bin or subset; simple*M*: The simple*M* procedure proposed by Gao et al. (2008) with *C* = 0.995. Results are averaged over 5000 simulations.

## 4 Application to colorectal cancer

We applied our proposed two-step approach to a genome-wide scan of gene-by-sex (G×Sex) interaction on the risk of colorectal cancer (CRC). The FIGI (Functionally Informed Gene-Environment Interaction) study is a multi-institutional collaborative effort to identify novel *G* × *E* interactions for CRC. Details of the FIGI study have been previously published (Schumacher et al., 2015; Schmit et al., 2019; Huyghe et al., 2019). In brief, epidemiological and genotype data were pooled from 61 cohort and case-control CRC studies from 3 large consortia - the Genetics and Epidemiology of Colorectal Cancer Consortium (GECCO), the Colorectal Transdisciplinary Study (CORECT), and Colon Cancer Family Registry (CCFR). Analyses include only individuals with complete exposure and covariate information, and were limited to individuals of European ancestry as determined by self-reported race and clustering of principal components with 1000 Genomes EUR sample. Details on genotyping, quality control, data collection and harmonization have been described previously (Hutter et al., 2012; Huyghe et al., 2019). Our analysis included *N* = 89, 304 individuals (40,647 cases and 48,657 controls) and *M* = 7, 809, 725 imputed SNPs. Autosomal SNPs were imputed to the Haplotype Reference Consortium r1.1 reference panel via the Michigan Imputation Server (Das et al., 2016). Imputed SNPs were filtered based on a pooled MAF ≥ 1% and imputation accuracy of *R*^2^ > 0.80.

Logistic regression models were used to model SNP allelic dosages (*G*) and sex (*E*) on case/control status (*Y*) genome-wide. The marginal outcome-gene association, regressing *Y* on *G* genome-wide, was used as the screening statistic in Step 1. All models included the following set of adjustment covariates - age, study/genotyping batch, and ancestry as defined by the first three principal components. Analyses using similar adjustment covariates were used in a recent investigation of G× Alcohol interaction in these data Jordahl et al. (2022).

Similar to the simulation study, bin significance threshold cutoffs for SB-weighted testing were based on cutoffs *τ* = (0, 5/*M*, 15/*M*, 35/*M*,…, 1). Bin flooding is apparent as bin 1 contains 3,795 SNPs (Figure 4) with a direct G-CRC association *p*-value < 5/7, 809, 725. Under the RB-weighted testing approach with *B*_0_ = 5, these 3,759 SNPs fill the first 8 bins. For the SB approach, the estimated effective number of independent tests among the 3,759 SNPs is *M*^*^ = 423 based on the simple*M* approach. Thus, each of the 3,795 *G* × *E* tests in bin 1 are evaluated against a significanceare tested against a significance threshold of 0.025/423. One SNP (18:46458950:G:A) surpassed the threshold in bin 1. This SNP is located near the SMAD7 gene, which has been previously shown as a potential marker of colorectal cancer (Thompson et al., 2009; Jiang et al., 2013; Stolfi et al., 2014; Alidoust et al., 2022). No statistically significant *G* × *E* effects were discovered using either the standard GWIS or the RB-weighted two-step approach.

**Figure 4:**
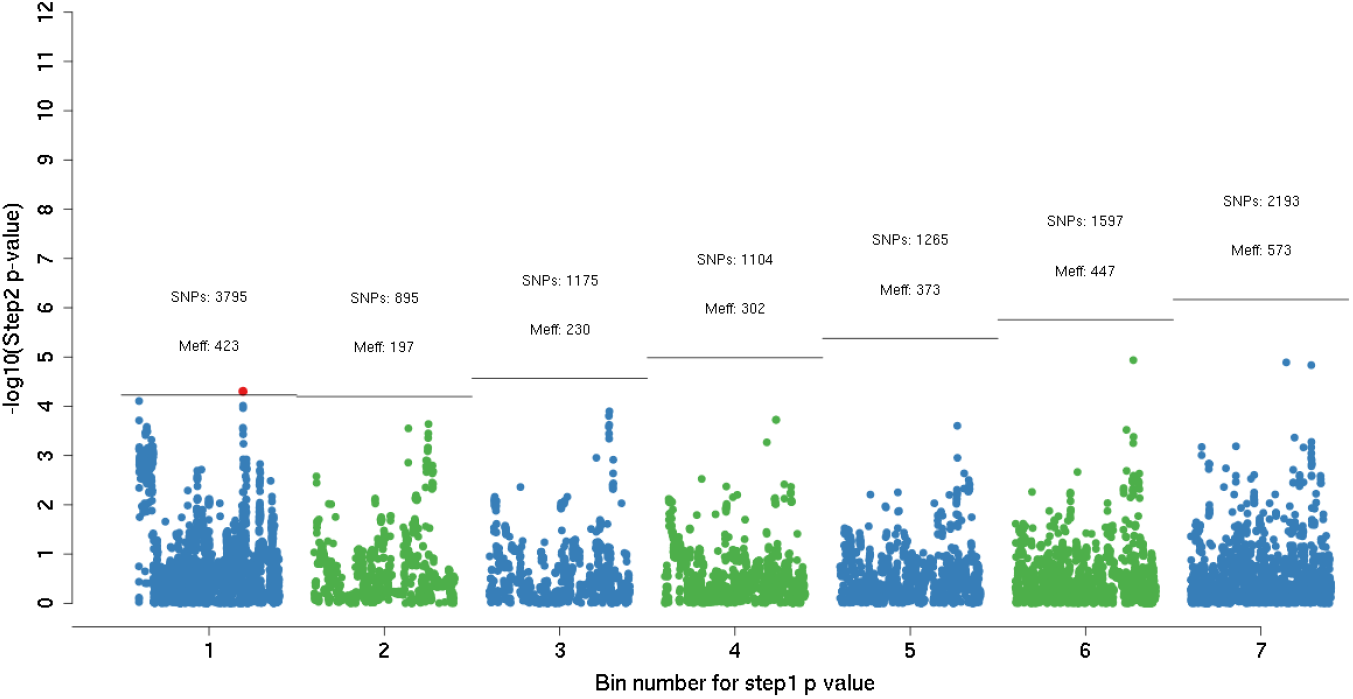
Results from the *G*-by-sex interaction scan using data provided by the FIGI consortium (*N* = 89, 304, *M* = 7, 809, 725). x-axis: Bins are based on the marginal outcome-gene association statistic (e.g. SNPs that have a Step 1 statistic < 5/*M* are included in bin 1). y-axis: *p*-value of the *G* × *E* association provided by the GWIS (on the – log_10_ scale). Number of SNPs in each bin as well as the effective number of independent SNPs using the simple*M* approach are included. Horizontal line indicates the threshold the Step 2 *p*-value must cross to be statistically significant, maintaining the overall FWER=0.05. Only SNPs in the first 7 bins are shown in this figure.

## 5 Discussion

Two-step tests generally provide greater power than standard one-step approaches for genome-wide *G* × *E* discovery (Gauderman et al., 2017). We propose a novel approach, significance-based (SB) weighted hypothesis testing, that aims to address the shortcomings of its predecessors. In Step 1, we allocate SNPs into bins based on the significance of the screening statistic rather than on their ranking. Then, in step 2, to account for SNP-SNP correlations due to LD, we control the bin-wise error rate based on an estimate of the effective number of independent tests. We show that SB-weighted testing is comparable to RB-weighted hypothesis testing when 1) no marginal *G*-only effects are present or 2) weak marginal *G*-only effect are present (i.e. little-to-no skew in the *p*-value distribution) and outperforms RB-weighted hypothesis testing when *G*-only loci flooding of the top bins exists. In addition, using an estimate of the effective number tests (simple*M*) to account for LD-based correlation among SNPs provides additional power for either of the two-step methods. We demonstrated our SB-weighted approach to identify novel gene-sex interactions for colorectal cancer using data provided by the FIGI consortium.

In our simulation study, we also show that power for the RB-weighted hypothesis test is sensitive to both the number and magnitude of the step 1 screening statistics as well as the initial bin size. This corresponds to the number of marginal genetic effects that are associated with the outcome if the outcome-gene association statistic proposed by Kooperberg and LeBlanc (2008) is used in Step 1. These factors, among others, will negatively affect the prioritization of the causal (*G* × *E*) loci in the testing step since the number of tests per bin are capped *a priori*. Alternatively, the use of significance cutoffs in the SB-weighted approach is robust to both the number and magnitude of the step 1 statistics and has greater power over RB-weighted testing when there is competition in bin prioritization. Adjusting for the effective number of tests using the simple*M* method of Gao et al. (2008) yields additional consistent, improvements in power while preserving the FWER.

Our two-step hypothesis testing procedure identified a new locus within SMAD7 that interacts with biological sex to modulate CRC risk. Further studies should examine the potential mechanisms through which this newly discovered locus impacts CRC risk. We note that *G*×Sex interaction for this locus would not have been found using either the standard one-step GWIS or the RB-weighted testing approach.

We envision several directions to further explore SB-weighted testing. First, the marginal outcome-gene association was used as the screening statistic in Step 1. However, SB-weighted testing should be valid using any screening statistic in Step 1 as long as it is independent of the testing statistic in Step 2 (Dai et al., 2012). Further investigation into the performance of SB-weighted testing is warranted under different phenotypes, sampling designs, and screening statistics. The binning of tests was performed by developing significance-level based cutoffs of the *p*-values in Step 1. Our rationale for this approach was to, in expectation, develop bins that were similar to the rank-based binnings proposed by Ionita-Laza et al. (2007). However, one can bin tests arbitrarily based on the *p*-value distribution of the Step 1 screening statistics or on the distribution of the screening statistics themselves. To this end, we have not yet considered alternative approaches for determining the cutoffs used for binning tests.

LD was taken into account through estimating the effective number of independent tests. While we explored the performance of the simple*M* procedure proposed by Gao et al. (2008), several other estimators have been suggested (Cheverud, 2001; Li and Ji, 2005; Galwey, 2009; Nyholt, 2004; Li et al., 2012). Alternative approaches to adjust *p*-values within bins (or the testing subset) can be adopted (Conneely and Boehnke, 2007). While genetic correlation within bins have been considered, between bin LD can be present as well. Methods to account for both within- and between-bin genetic correlation may further improve power and should be investigated further. Alternatively, one may instead control the false discovery rate (Benjamini and Hochberg, 1995), the expected proportion of discoveries that are false, rather than the FWER. Future research is needed into developing FDR-based two-step hypothesis tests.

We have demonstrated that the current two-step *G* × *E* testing paradigm can be greatly improved by 1) incorporating a more robust binning procedure for the screening step (Step 1) and 2) taking into account LD when adjusting the BWER. Our proposal suggests to bin test using a significance-based weighting scheme and to adjust the BWER by using the simple*M* procedure. We have demonstrated, by simulation and application to a colorectal cancer study, that these modifications have the potential to discover novel *G*×*E* interactions for a complex trait. Additional examinations of the binning approach and the method to account for LD to further improve 2-step GWIS power are warranted.

## Acknowledgements

The work was partially supported by NIH grants T32ES013678, P01CA196569, R01CA201407, and P30ES007048. These awards had no influence over the experimental design, data analysis or interpretation, or writing the manuscript.

## S1 Appendix: Additional figures and tables

**Figure S1:**
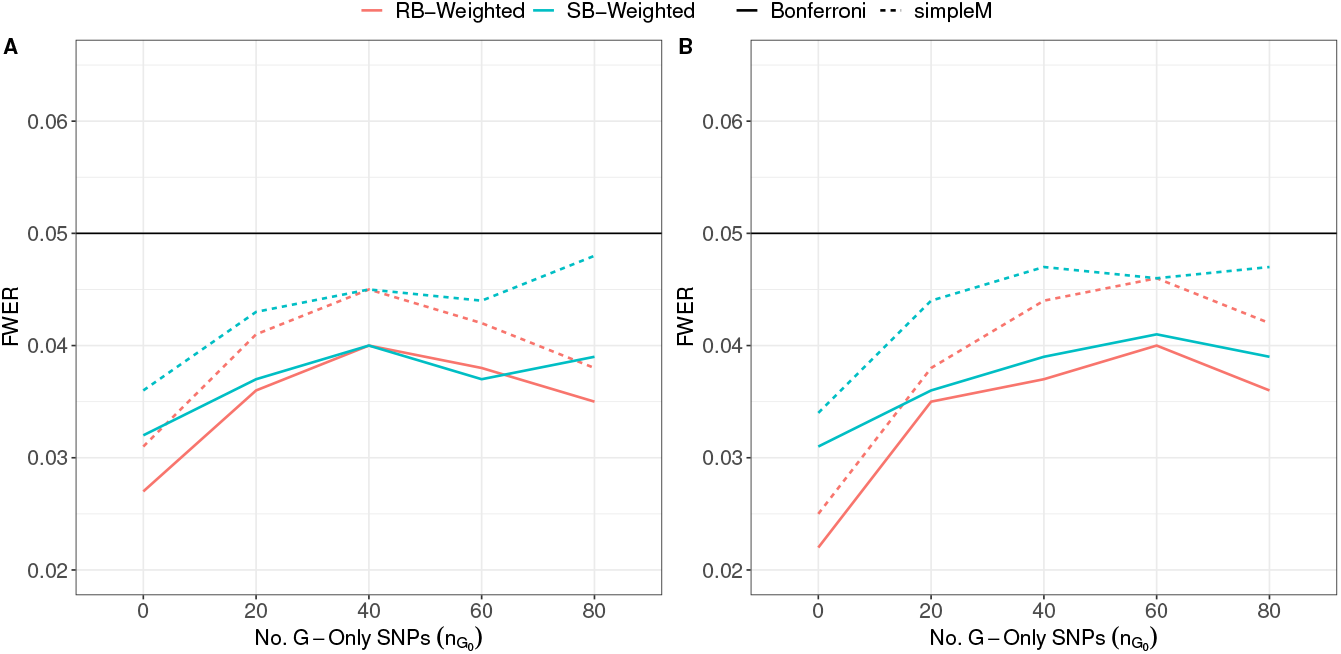
Estimated FWER when *n_G_ G*-only effects are present (*n_G_* ∈ {10, 20, 40, 80}) each explaining 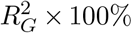 of the variation in the quantitative trait. RB-Weighted: Rank-based two-step weighted testing with initial bin size 5 in Step 1; SB-Weighted: Significance-based weighted hypothesis testing with *τ* = (0, 5/25000, 15/25000,…, 1) as the *p*-value cutoffs in Step 1. Bonferroni: Standard Bonferroni correction within bin; simple*M*: The simple*M* procedure proposed by Gao et al. (2008) with *C* = 0.995. Results are averaged over 5000 simulations. Panel A: 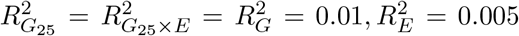, *N* = 2, 000; Panel B: 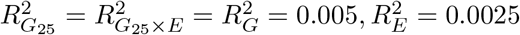, *N* = 4, 000.

**Figure S2:**
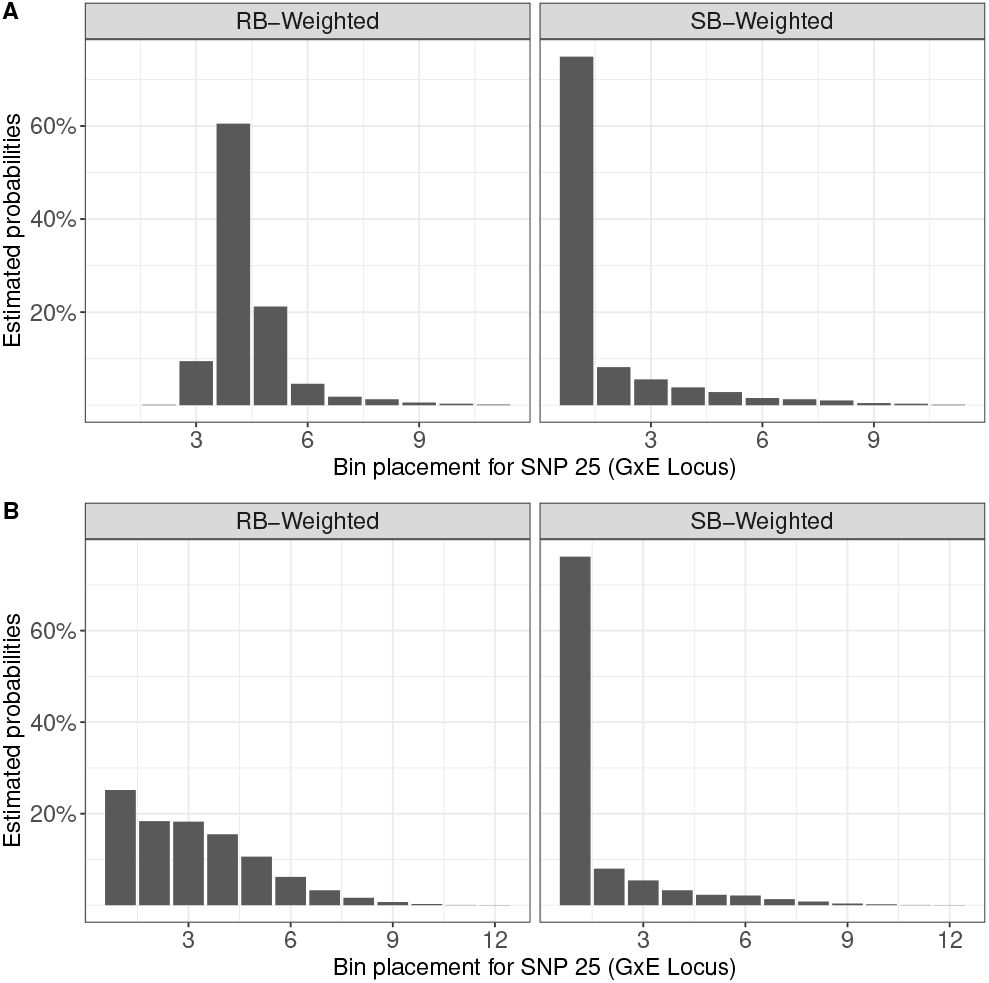
Bar chart of bin placement for the 25th SNP (i.e. *G* × *E* locus) in Step 1 over 5,000 simulations. RB-Weighted: Rank-based weighted hypothesis testing using with initial bin size *B*_0_ in Step 1; SB-Weighted: Significance-based weighted hypothesis testing using *τ* = (0, *B*_0_/25000, 3*B*_0_/25000,…, 1) as the *p*-value cutoffs in Step 1. Simulation parameters: 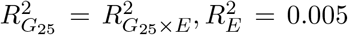, *N* = 2, 000, *M* = 25, 000. Panel A) *n_G_* = 10 *G*-only SNPs each with 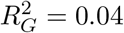; Panel B: *n_G_* = 80 *G*-only SNPs each with 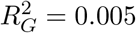.

**Table S1:** Estimated FWER when *n_G_ G*-only effects are present (*n_G_* ∈ {10, 20, 40, 80} and the total amount of variation explained is fixed at 40% 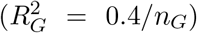. RB Weighted: Rank-based weighted hypothesis testing proposed by Ionita-Laza et al. (2007) with *B*_0_ = 5; SB-Weighted: Our proposed significance-based weighted hypothesis testing with *τ* = (0, 5/25000, 15/25000,…, 1) as the *p*-value cutoffs. Bonferroni: Standard Bonferroni correction within bin; simple*M*: The simple*M* procedure proposed by Gao et al. (2008) with *C* = 0.995. Results are averaged over 5000 simulations.

|  |  |  |  |  |
| --- | --- | --- | --- | --- |
| $n_G =$ | 10 | 20 | 40 | 80 |
| $R_G^2 =$ | 0.04 | 0.02 | 0.01 | 0.005 |
| **RB-Weighted** |  |  |  |  |
| Bonferroni | 0.044 | 0.040 | 0.035 | 0.036 |
| simpleM | 0.045 | 0.045 | 0.039 | 0.039 |
| **SB-Weighted** |  |  |  |  |
| Bonferroni | 0.037 | 0.037 | 0.036 | 0.036 |
| simpleM | 0.045 | 0.045 | 0.043 | 0.044 |

## S2 Appendix: Simulation setup

Let **G** be an *N* × *M* genotype matrix for *N* individuals and *M* SNPs. We partition the *M* SNPs into blocks of 50 SNPs such that **G** = [**G**_1_, **G**_2_,…] where **G***_j_* is the *j*th block of *N* × 50 SNPs. Each **G***_j_* is simulated based on sampled minor allele frequencies (MAFs) and LD-matrices from the 1000 Genomes Project. For clarity, we denote *G_j_* as the *j*th SNP and **G***_j_* as the *j*th block. Quantitative traits are simulated according to the following linear model:

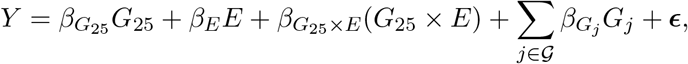

where 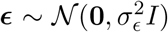 for some 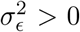, *E* is the exposure variable (assumed to be binary) with Pr(E = 1) = 0.3 and 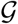 corresponds to the set of SNPs that are only marginally associated with the outcome but have no *G* × *E* effect (G-only loci). By construction, the 25th SNP within block 1 (**G**_1_) has a true *G* × *E* effect on the outcome (i.e. the *G* × *E* locus). 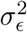 was chosen to explain the remaining variance in the outcome after accounting for the variance explained by the causal *G* × *E* locus, the exposure, and the G-only loci across the different scenarios outlined in the main text.

The *M* = 25, 0000 genotypes are simulated in blocks (**G** = [**G**_1_, **G**_2_,…, **G**_500_]) such that each block consists of 50 SNPs drawn from a mean zero multivariate normal distribution with variance-covariance matrix based on the LD pattern derived from a sampled region of the 1000 Genomes Project. The normal variates are then trichotomized into genotypes based on the 1000 Genomes Project derived MAFs assuming Hardy-Weinberg equilibrium. Thus, genotypes are correlated within a block but independent across blocks. Define **V** = [**V**_1_,…, **V**_500_] and **f** = (**f**_1_,…, **f**_500_), where **V***_j_* is a 50 × 50 LD matrix and *f_j_* is the corresponding vector of minor allele frequencies (MAF) of the 50 SNPs for *j* = 1,…, 500. Both **V***_j_* and *f_j_* are derived from a randomly sampled region from the 1000 Genomes Project. To avoid storing 500 unique values of **V***_j_* and *f_j_*, we only store 50 unique values (randomly sampled regions), and recycled them such that the (**V**_1_, **f**_1_) = (**V**_51_, **f**_51_) = (**V**_101_, **f**_101_),…, (**V**_2_, **f**_2_) = (**V**_52_, **f**_52_) = (**V**_102_, **f**_102_),…, (**V**_3_, **f**_3_) = (**V**_53_, **f**_53_) = (**V**_103_, **f**_103_),….

To simulate allelic dosages, first let 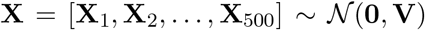 be a *N × M* matrix of mean zero normal variates with block correlation structure **V**. Letting *G_i,k_* and *X_i,k_* be the ith row of the kth column of **G** and **X** (*i* = 1,…, *N; k* = 1,…, *M*), respectively, and being the kth element of **f**,

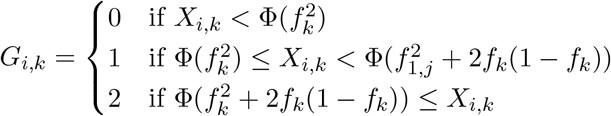

where Φ(.) is the cumulative distribution function of the standard normal distribution.

